# Feature Extraction Approaches for Biological Sequences: A Comparative Study of Mathematical Models

**DOI:** 10.1101/2020.06.08.140368

**Authors:** Robson Parmezan Bonidia, Lucas Dias Hiera Sampaio, Douglas Silva Domingues, Alexandre Rossi Paschoal, Fabrício Martins Lopes, André Carlos Ponce de Leon Ferreira de Carvalho, Danilo Sipoli Sanches

## Abstract

The number of available biological sequences has increased significantly in recent years due to various genomic sequencing projects, creating a huge volume of data. Consequently, new computational methods are needed to analyze and extract information from these sequences. Machine learning methods have shown broad applicability in computational biology and bioinformatics. The utilization of machine learning methods has helped to extract relevant information from various biological datasets. However, there are still several obstacles that motivate new algorithms and pipeline proposals, mainly involving feature extraction problems, in which extracting significant discriminatory information from a biological set is challenging. Considering this, our work proposes to study and analyze a feature extraction pipeline based on mathematical models (Numerical Mapping, Fourier, Entropy, and Complex Networks). As a case study, we analyze Long Non-Coding RNA sequences. Moreover, we divided this work into two studies, e.g., (I) we assessed our proposal with the most addressed problem in our review, e.g., lncRNA vs. mRNA; (II) we tested its generalization on different classification problems, e.g., circRNA vs. lncRNA. The experimental results demonstrated three main contributions: (1) An in-depth study of several mathematical models; (2) a new feature extraction pipeline and (3) its generalization and robustness for distinct biological sequence classification.

## 1. Background

In recent years, due to advances in DNA sequencing, an increasing number of biological sequences have been generated by thousands of sequencing projects [1], creating a huge volume of data [2]. During the last decade, Machine Learning (ML) methods have shown broad applicability in computational biology and bioinformatics [3]. Consequently, the ability to process and analyze biological data has advanced significantly [4]. Tools have been applied in gene networks, protein structure prediction, genomics, proteomics, protein-coding genes detection, disease diagnosis, and drug planning [5, 6]. Fundamentally, ML investigates how computers can learn (or improve their performance) based on the data. Moreover, ML is a specialization of computer science related to pattern recognition and artificial intelligence [7].

Based on this, several works have focused on investigating sequences of DNA and RNA molecules [8, 9, 10]. Applying ML methods in these sequences has helped to extract important information from various datasets to explain biological phenomena [3]. The development of efficient approaches benefits the mathematical understanding of the structure of biological sequences [1], e.g., Precision cancer diagnostics [11], analytics in plants [12], and Coronavirus epidemic [13, 14]. However, according to [3, 15], there are still several challenging biological problems that motivated the emergence of proposals for new algorithms. Fundamentally, biological sequence analysis with ML presents one major problem, e.g., Feature Extraction [16], an inevitable process, especially in the stage of biological sequence preprocessing [10, 17].

Feature extraction seeks to generate a feature vector, optimally transforming the input data [16]. This procedure is exceptionally relevant for the success of the ML application because another primary goal is to extract important information from input data compactly, as well as removing noise and redundancy to increase the accuracy of ML models [18, 16]. Necessarily, several methods in bioinformatics apply ML algorithms for sequence classification, and as many algorithms can deal only with numerical data, sequences need to be translated into sequences of numbers.

Thereby, modern applications extract relevant features from sequences based on several biological properties, e.g., physicochemical, Open Reading Frames (ORF)-based, usage frequency of adjoining nucleotide triplets, GC content, among others. This approach is common in biological problems, but these implementations are often difficult to reuse or adapt to another specific problem, e.g., ORF features are an essential guideline for distinguishing Long non-coding RNAs (lncRNA) from protein-coding genes [19], but not useful features for classifying lncRNA classes [20, 21] (e.g., in [21], ORF score (feature importance) is less than 0.009 to classify circular RNA from other types of lncRNAs). Consequently, the feature extraction problem arises, in which extracting a set of useful features that contain significant discriminatory information becomes a fundamental step in the construction of a predictive model [22].

Therefore, these problems make the process of biological sequence classification a challenging task, creating a growing need to develop new techniques and methods to analyze sequences effectively and efficiently. Thereby, this work studies the performance of different feature extraction methods for biological sequence analysis, using mathematical models, e.g., numerical mapping, Fourier transform, entropy, and graphs. As a case study, we will use lncRNA sequences, which are fundamentally unable to produce proteins [23] and have recently casted doubt on its functionality [24].

LncRNAs present several problem classes (e.g., lncRNA vs. mRNA [25, 26] and lncRNA vs. circRNA [27]), thus enabling us to create a scenario to answer the questions raised in this work. Fundamentally, our main objective is to propose generalist techniques, demonstrating their efficiency concerning biological features. We consider biological approaches, those characteristics that present a bias to the analyzed problem or some biological explanation, e.g., ORF for lncRNA vs. mRNA [6, 19], as well as mathematical approaches and information quantity measures such as entropy. Based on this context and objectives, we assume the following hypothesis:

- **Hypothesis:** Feature extraction approaches based on mathematical models are as efficient and generalist as biological approaches.

Considering this, our work contributes to the area of computer science and bioinformatics. Specifically, it introduces new ideas and analysis for the feature extraction problem in biological sequences. Thereby, we present four new contributions: (1) A feature extraction pipeline using mathematical models; (2) Analysis of 9 mathematical models; (3) Analysis of 6 numerical mappings with Fourier, proposing statistical characteristics; (4) The generalization and robustness of mathematical approaches for the feature extraction in biological sequences.

## 2. Related Works

Essentially, as emphasized, we adopt lncRNA sequences as a case study, a class of Non-Coding RNAs (ncRNAs). Fundamentally, ncRNAs are unable to produce proteins. However, these ncRNAs contain unique information that produces other functional RNA molecules [28, 23]. Moreover, they demonstrate essential roles in cellular mechanisms, playing regulatory roles in a wide variety of biological reactions and processes [29, 28]. The ncRNAs can be classified by length into two classes: Long Non-Coding RNA (lncRNA - 200 nucleotides (nt) or more) and short ncRNA (less than 200 nt) [30, 31]. The lncRNAs are sequences with a length greater than 200 nucleotides [32], and according to recent studies, play essential roles in several critical biological processes [33, 34, 35], including transcriptional regulation [36], epigenetics [37], cellular differentiation [38], and immune response [39]. Moreover, they are correlated with some complex human diseases, such as cancer and neurodegenerative diseases [6, 40, 41].

In plants, according to [6, 42], the lncRNAs act in gene silencing, flowering time control, organogenesis in roots, photomorphogenesis in seedlings, stress responses [43, 44], and reproduction [45]. Furthermore, lncRNAs are present in large numbers in genome [46] and have similar sequence characteristics with protein-coding genes, such as 5’ cap, alternative splicing, two or more exons [47], and polyA+ tails [48]. They are also observed in almost all living beings, not only in animals and plants but also yeasts, prokaryotes, and even viruses [49, 50].

According to [46], lncRNAs do not contain functional ORFs. However, recent studies have found bifunctional RNAs [51], raising the possibility that many protein-coding genes may also have non-coding functions. Furthermore, lncRNAs can be grouped into five broad categories. The classification occurs conforming to the genomic location, that is, where they are transcribed, concerning well-established markers, e.g., protein-coding genes. Among the categories are [52, 47]: sense, antisense, bidirectional, intronic, intergenic. The genomic context does not necessarily provide some information about the lncRNAs function or evolutionary origin; nevertheless, it can be used to organize these broad categories [53].

In this context, we have conducted an in-depth review of the lncRNAs classification methods, in which several approaches have been developed, such as: CPC [54], CPAT [55], CNCI [56], PLEK [57], lncRNA-MFDL [58], LncRNA-ID [59], lncRScan-SVM [60], LncRNApred [61], DeepLNC [62], PlantRNA Sniffer [63], PLncPRO [64], RNAplonc [65], BASiNET [66], LncFinder [26], CREMA [67], LncRNAnet [19], CNIT [68], PLIT [69], PredLnc-GFStack [70], LGC [71] and DeepCPP [72]. For better understanding, Figure 1 presents theses works divided into Mathematical, Biological, and Hybrid approaches.

**Figure 1:**
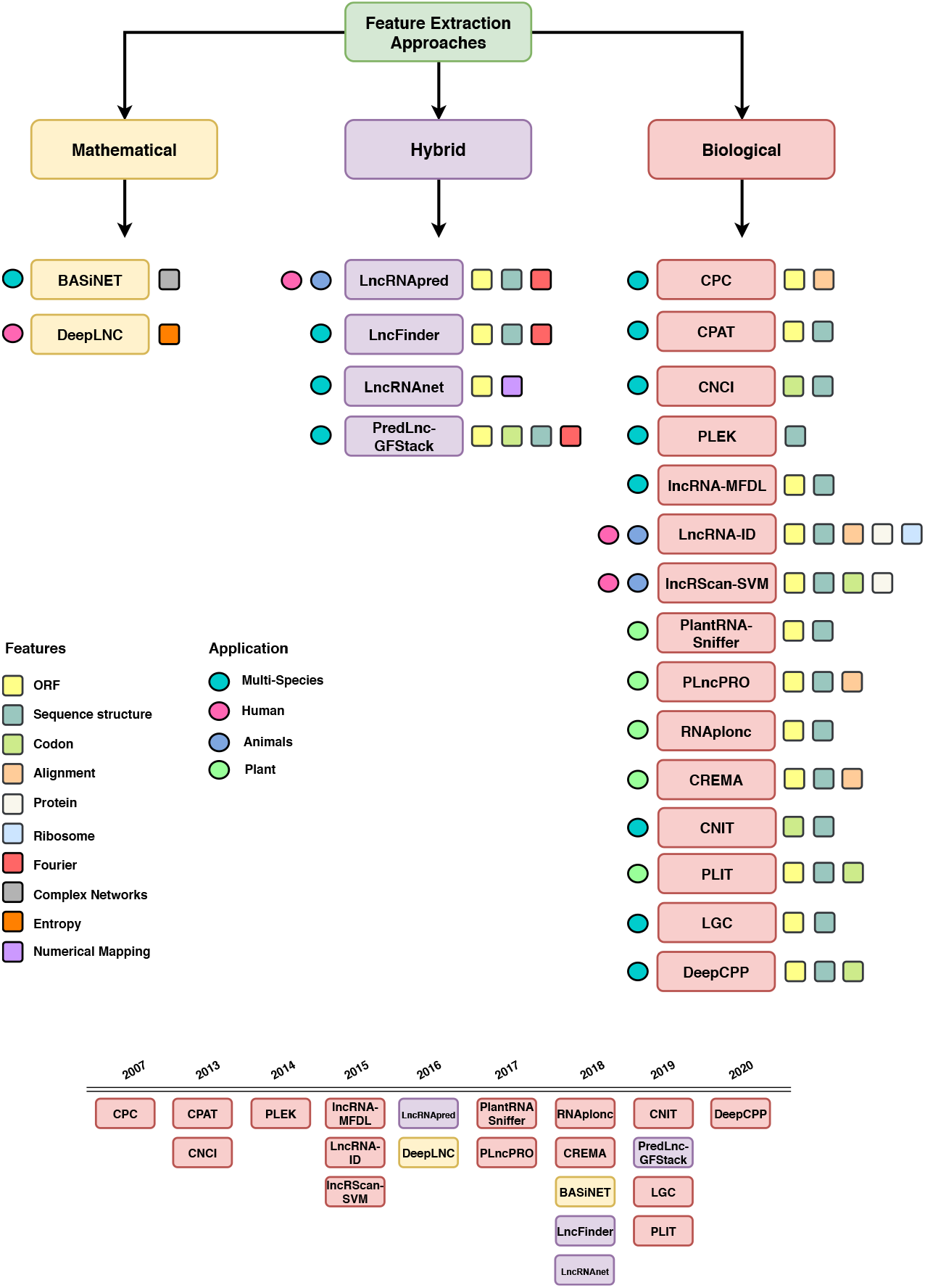
Feature extraction approaches in our case study divided into: Mathematical, Biological, and Hybrid.

The CPC uses the extent and quality of the ORF, and derivation of the BLASTX [73] search to measure the protein-coding potential of a transcript. In the classification, the authors applied the LIBSVM package to train a Support Vector Machine (SVM) model, using the standard radial basis function kernel. CPAT classifies transcripts of coding and non-coding using the Logistic Regression (LR) classifier. This approach implements four features: ORF coverage, ORF size, hexamer usage bias, and Fickett TESTCODE statistic. CNCI was induced with SVM and applies profiling Adjoining Nucleotide Triplets, and most-like CDS (MLCDS).

In contrast, PLEK (2014) is based on the k-mer scheme (*k* = 1, …, 5) to predict lncRNA, also applying the SVM classifier. lncRNA-MFDL uses Deep Learning (DL) and multiple features, among them: ORF, K-mer (k = 1, 2, 3), secondary structure (minimum free energy), and MLCDS. LncRNA-ID predicts lncRNAs with Random Forest (RF) through ORF (length and coverage), sequence structure (Kozak motif), ribosome interaction, alignment (profile Hidden Markov Mode - profile HMM), and protein conservation.

lncRScan-SVM uses stop codon count, GC content, ORF (score, CDS length and CDS percentage), transcript length, exon count, exon length, and average PhastCons scores. LncRNApred classified lncRNAs with RF and features based on ORF, signal to noise ratio, k-mer (k = 1, 2, 3), sequence length, and GC content. DeepLNC uses only the k-mer scheme with entropy and Deep Neural Network (DNN). PlantRNA Sniffer was developed in 2017 to predict Long Intergenic Non-Coding RNAs (lincRNAs). The method applied SVM and extracted features from ORF (proportion and length) and nucleotide patterns.

PLncPRO is based on machine learning and uses RF. The features selected include ORF quality (score and coverage), number of hits, significance score, total bit score, and frame entropy. RNAplonc classified sequences with the REPtree algorithm, considering 16 features (ORF, GC content, K-mer scheme (*k* = 1, …, 6), sequence length). BASiNET classifies sequences based on the feature extraction from complex network measurements. LncFinder tests five classifiers (LR, SVM, RF, Extreme Learning Machine, and Deep Learning), to apply the algorithm that obtains the highest accuracy. The authors extract features from ORF, secondary structural, and EIIP-based physicochemical properties.

CREMA uses ensemble machine learning classifiers. Features include mRNA length, ORF (length), GC content, Fickett score, hexamer score, alignment, transposable element, and sequence percent divergence from a transposable element. LncRNAnet applies a deep learning-based approach using numerical mapping and ORF indicators. CNIT is the updated CNCI tool with a novel approach (XGBoost models with adjoining nucleotide triplets and MLCDS). PLIT is a new alignment-free tool that uses ORF, transcript length, Fickett score, Hexamer Score, GC content, and codon-bias features.

Lastly, PredLnc-GFStack also uses the stacked ensemble learning method by extracting features based on codon-bias, Fickett score, ORF, GC content, coding sequence, transcript length, k-mer, CTD, Hexamer score, signal to noise ratio, UTR coverage, EDP of transcripts (entropy density profiles) and structure-related. LGC proposes a feature relationship-based approach (ORF length and GC content). DeepCPP is a deep learning method for RNA coding potential prediction. Among the extracted features are ORF, hexamer score, Fickett score, k-mer, g-gap, and nucleotide bias.

In general, the aforementioned works apply supervised learning methods using binary classification (two classes - lncRNAs and protein-coding genes (mRNA)). There is a considerable amount of research on humans, followed by animals and plants. Regarding feature extraction, we observed a full domain of ORF and sequence-structure descriptors. As seen in Figure 1, there is a frequent use of biological features. On the other hand, some works have explored mathematical approaches for feature extraction, such as Genomic Signal Processing (GSP), DNA Numerical Representation (DNR) [61, 26], and Complex Networks [66]. Nevertheless, the authors used these characteristics in conjunction with other biological feature extraction techniques or without testing other mathematical features. Practically no papers have focused on several mathematical approaches. Based on this, the objective of this section was to summarize the main methods of the literature and their characteristic descriptors. Therefore, we will not use the works shown for comparison, but the most applied features.

## 3. Materials and Methods

In this section. we describe the methodological approach used to achieve the proposed objectives, as shown in Figure 2. Essentially, we divided our study into five stages: (1) Data selection and preprocessing; (2) Feature extraction; (3) Training; (4) Testing; (5) Performance analysis. Hence, each stage of the study is described, as well as information about the adopted process.

**Figure 2:**
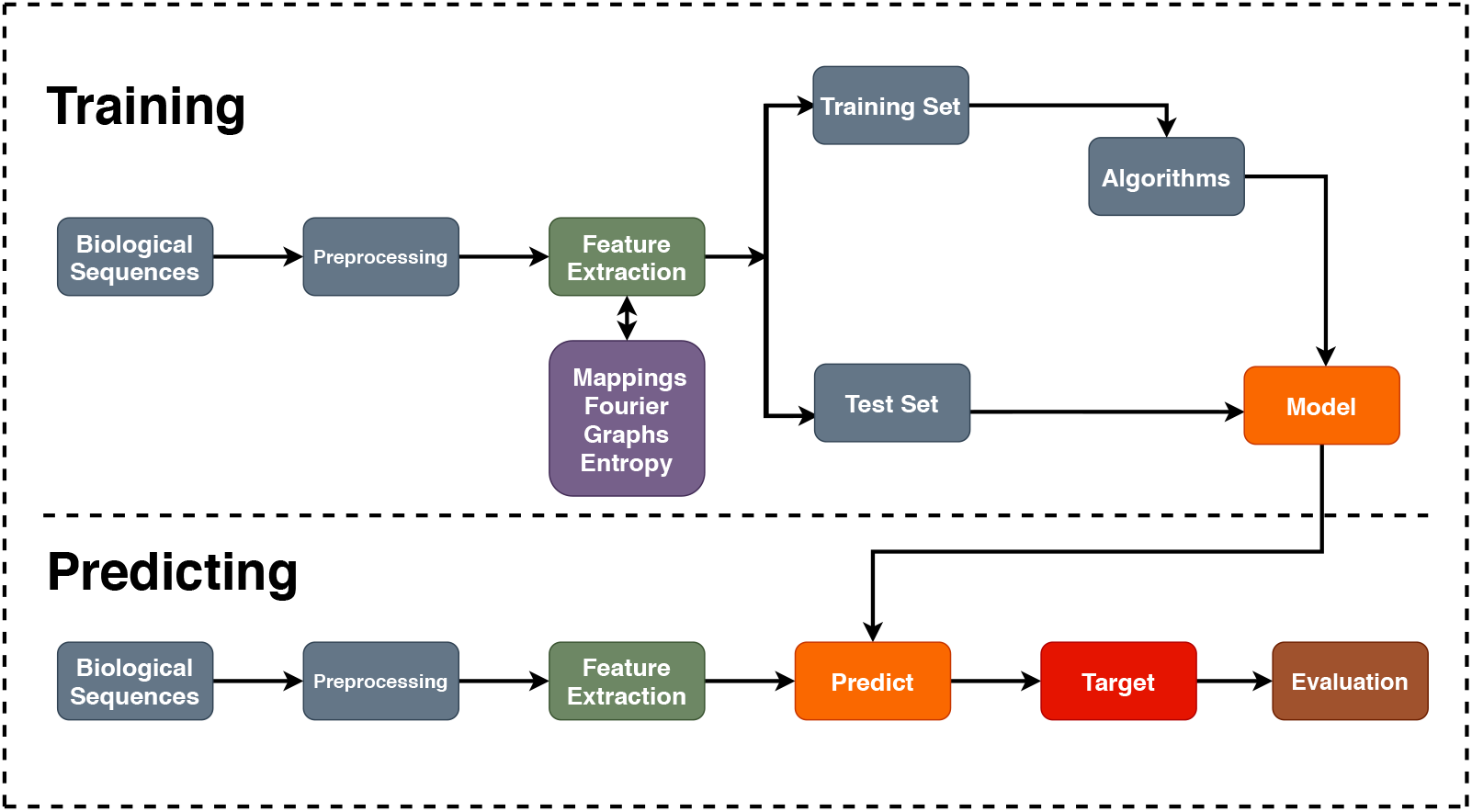
Proposed Pipeline. Essentially, (1) datasets are preprocessed; (2) Feature extraction techniques are applied to each dataset; (3) Machine learning algorithms are executed in the training set to induce predictive models; (4) Induced models are applied to the test set; Finally, (5) the models are evaluated.

This work was also divided into two case studies: (I) We assessed our mathematical approaches with the most addressed problem in our review, e.g., lncRNA vs. mRNA; (II) We tested its generalization on different classification problems.

### 3.1. Data Selection

As previously mentioned, we chose the lncRNAs classification problem, because it is a new and relevant theme in the literature, in which, recently, it has presented several works, mainly with ML, as explored in Section 2. However, we will also adopt other datasets to assess the generalization of mathematical features. As preprocessing, we used only sequences longer than 200*nt* [57], and we also removed sequence redundancy. Moreover, the sampling method was adopted in our dataset, since we are faced with the *imbalanced data problem* [20]. Therefore, we applied random majority under-sampling, which consists of removing samples from the majority class (to adjust the class distribution) [74]. Finally, we divided this paper into two case studies.

#### 3.1.1. Case Study I

Sequences of five plant species were adopted to validate the proposed approaches. The summary of the dataset can be seen in Table 1. According to the literature approaches, this study also adopts two classes for the datasets: the positive class, with lncRNAs, and the negative class, with protein-coding genes (mRNAs).

**Table 1:** Adopted species to create the datasets.

| Species | Sequences | Samples | Preprocessing | Selected |
| --- | --- | --- | --- | --- |
| *A. trichopoda* | lncRNA | 5698 | 4556 | 4556 |
|  | mRNA | 26846 | 22326 | 4556 |
| *A. thaliana* | lncRNA | 2540 | 2540 | 2540 |
|  | mRNA | 13973 | 13973 | 2540 |
| *C. sinensis* | lncRNA | 2562 | 2215 | 2215 |
|  | mRNA | 46147 | 45846 | 2215 |
| *C. sativus* | lncRNA | 1929 | 1730 | 1730 |
|  | mRNA | 30364 | 29829 | 1730 |
| *R. communis* | lncRNA | 4198 | 3487 | 3487 |
|  | mRNA | 31221 | 29042 | 3487 |

The mRNA data of the *Arabidopsis thaliana* (obtained from CPC2 [25]) were built from the `RefSeq database` with protein sequences annotated by Swiss-Prot [25], and lncRNA data from the `Ensembl` (*v*87) and `Ensembl Plants` (*v*32) database. The mRNA transcript data of the *Amborella trichopoda*, *Citrus sinensis*, *Cucumis sativus* and *Ricinus communis* were extracted from `Phytozome` (version 13) [75]. The lncRNAs data from these species were extracted from `GreeNC` (version 1.12) [76].

#### 3.1.2 Case Study II

In this case study, we will apply the best mathematical models (considering accuracy) of case study I to different classification problems with lncRNAs, in order to test their generalization. Thus, divided this part into three problems:

- **Problem 1** (lncRNA vs. sncRNA): Dataset with only non-coding sequences (lncRNA and Small non-coding RNAs (sncRNAs), also obtained from [25])

- lncRNA: 1291 sequences — sncRNA: 1291 sequences
- **Problem 2** (lncRNA vs. Antisense): Dataset with lncRNAs and long noncoding antisense transcripts (obtained from [77]).

- lncRNA: 57 sequences — Antisense: 57 sequences
- **Problem 3** (circRNA vs. lncRNA): Dataset with lncRNA and circular RNAs (cirRNAs) sequences (circRNA obtained from PlantcircBase [78]. This problem was based on [21] and [27], in order to classify circRNA from other lncRNAs.

- circRNA: 2540 sequences — lncRNA: 2540 sequences

It is important to emphasize that we used only sequences from *Arabidopsis thaliana* in this second case study because it is the model species in plants. Moreover, plant sequences is the least addressed field by the studies, consequently presenting more challenges.

### 3.2. Feature Extraction

In this section, 9 feature extraction approaches are shown: 6 numerical mapping techniques with Fourier transform, Entropy, Complex Networks. It is necessary to emphasize that we denote a biological sequence **s** = (*s*[0], *s*[1], …, *s*[*N* − 1]) such that **s** ∈ {*A, C, G, T*}^*N*^ [20].

### 3.3. Fourier Transform and Numerical Mappings

To extract features based on a Fourier model, we applied the Discrete Fourier Transform (DFT), widely used for digital image and signal processing (here GSP), which can reveal hidden periodicities after transformation of time domain data to frequency domain space [79]. According to Yin and Yau [80], the DFT of a signal with length *N*, **x** ∈ ℝ^*N*^, at frequency *k*, can be defined by Equation (1):

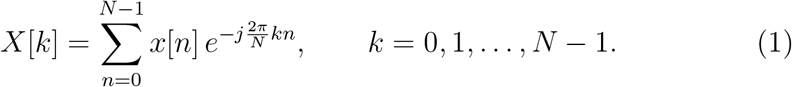

This method is has been widely studied in bioinformatics, mainly for analysis of periodicities and repetitive elements in DNA sequences [81] and protein structures [82]. This approach is shown in Figure 3 and was based on [20].

**Figure 3:**
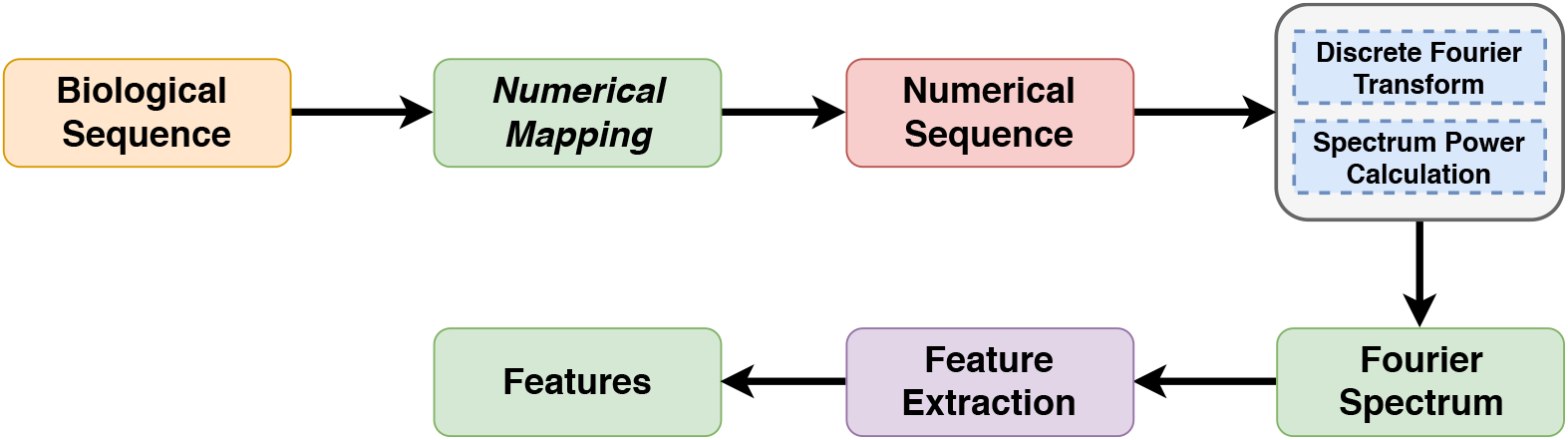
Fourier Transform and Numerical Mapping Pipeline. (1) Each sequence is mapped to a numerical sequence; (2) DFT is applied to the generated sequence; (3) The spectrum power is calculated; (4) The Feature Extraction is performed; Finally, (5) the features are generated.

To calculate DFT, we will use the Fast Fourier Transform (FFT), that is a highly efficient procedure for computing the DFT of a time series [83]. However, to use GSP techniques, a numeric representation should be used for the transformation or mapping of genomic data. In the literature, distinct DNR techniques have been developed [84]. According to MendizabalRuiz et al. [85], these representations can be divided into three categories: single-value mapping, multidimensional sequence mapping, and cumulative sequence mapping. Thereby, we study 6 numerical mapping techniques (or representations), which will be presented below: Voss [86], Integer [85, 87], Real [88], Z-curve [89], EIIP [90] and Complex Numbers [84, 91, 92].

#### 3.3.1. Voss Representation

This representation can use single or multidimensional vectors. Fundamentally, this approach transforms a sequence **s** ∈ {*A, C, G, T*}^*N*^ into a matrix **V** ∈ {0, 1}^4×*N*^ such that **V** = [**v**_1_, **v**_2_, **v**_3_, **v**_4_]^*T*^, where *T* is the transpose operator and each **v***_i_* array is constructed according to the following relation

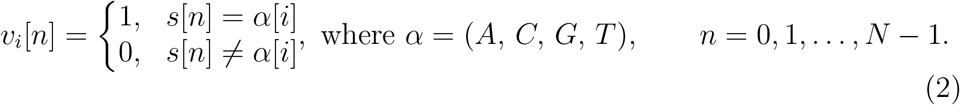

As a result, each row of matrix **V** may be seen as an array that marks each base position such that the first row denotes the presence of base *A*, row two for base *C*, row three base *G* and the last row for base *T*. For example, let **s** = (*G, A, G, A, G, T, G, A, C, C, A*) be a sequence that needs to be represented using Voss representation, therefore, **v**_1_ = (0, 1, 0, 1, 0, 0, 0, 1, 0, 0, 1), which represents the locations of bases *A*, **v**_2_ = (0, 0, 0, 0, 0, 0, 0, 0, 1, 1, 0) for bases *C*, **v**_3_ = (1, 0, 1, 0, 1, 0, 1, 0, 0, 0, 0) for the *G* bases, **v**_4_ = (0, 0, 0, 0, 0, 1, 0, 0, 0, 0, 0) for *T* bases. Then, using the DFT in the indicator sequences shown above, we obtain (see Equation 3):

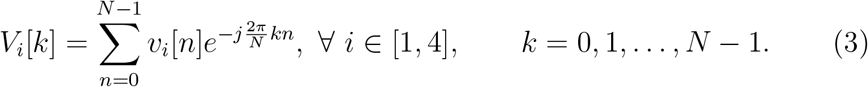

The power spectrum of a biological sequence can be obtained by Equation (4):

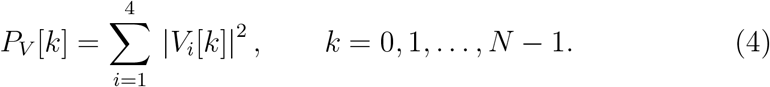

#### 3.3.2. Integer Representation

This representation is one-dimensional [87, 85]. This mapping can be obtained by substituting the four nucleotides (T, C, A, G) of a biological sequence for integers (0, 1, 2, 3), respectively, e.g., let **s** = (G, A, G, A, G, T, G, A, C, C, A), thus, **d** = (3, 2, 3, 2, 3, 0, 3, 2, 1, 1, 2), as exposed in Equation (5). The DFT and power spectrum are presented in Equation (6).

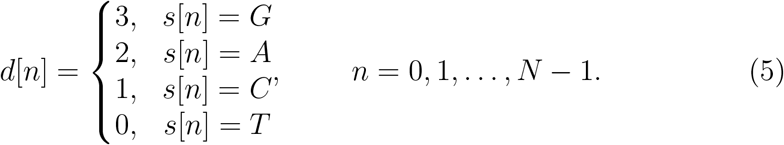

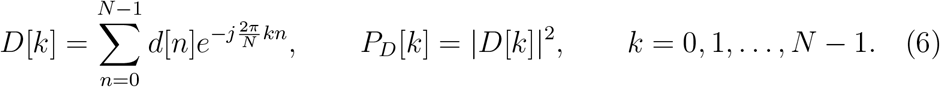

#### 3.3.3. Real Representation

In this representation, Chakravarthy et al. [88] use real mapping based on the complement property of the complex mapping of [81]. This mapping applies negative decimal values for the purines (*A, G*), and positive decimal values for the pyrimidines (*C, T*), e.g., let **s** = (*G, A, G, A, G, T, G, A, C, C, A*), thus, **r** = (−0.5, −1.5, −0.5, −1.5, −0.5, 1.5, −0.5, −1.5, 0.5, 0.5, −1.5), as Equation (7) and Equation (8).

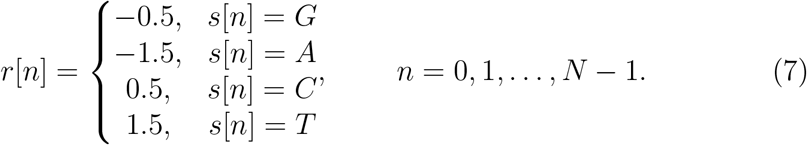

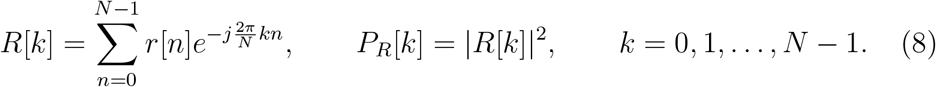

#### 3.3.4. Z-curve Representation

The Z-curve scheme is a three-dimensional curve presented by [89], to encode DNA sequences with more biological semantics. Essentially, we can inspect a given sequence *s*[*n*] of length *N*, taking into account the *n*-th element of the sequence (*n* = 1, 2, …, *N*). Then, we denote the cumulative occurrence numbers *A_n_*, *C_n_*, *G_n_* and *T_n_* for each base *A*, *C*, *G* and *T*, as the number of times that a base occurred from *s*[1] up until *s*[*n*]. Fundamentally, this method reduces the number of indicator sequences from four (Voss) to three (Z-curve) in a symmetrical way for all four components [93]. Therefore:

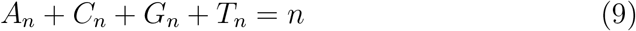

Where the Z-curve consists of a series of nodes *P*_1_, *P*_2_, …, *P_N_*, whose coordinates *x*[*n*], *y*[*n*], and *z*[*n*] (*n* = 1, 2, …, *N*) are uniquely determined by the Z-transform, shown in Equation (10):

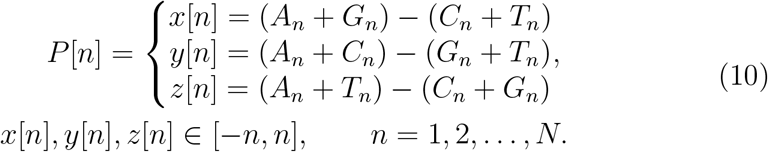

The coordinates *x*[*n*], *y*[*n*], and *z*[*n*] represent three independent distributions that fully describe a sequence [84]. Therefore, we will have three distributions with definite biological significance: (1) *x*[*n*] = purine/pyrimidine, (2) *y*[*n*] = amino/keto, (3) *z*[*n*] = weak hydrogen bonds/strong hydrogen bonds [89], e.g., let **s** = (G, A, G, A, G, T, G, A, C, C, A), thus, **x** = (1, 2, 3, 4, 5, 4, 5, 6, 5, 4, 5); **y** = (−1, 0, −1, 0, −1, −2, −3, −2, −1, 0, 1); **z** = (−1, 0, −1, 0, −1, 0, −1, 0, −1, −2, −1). Essentially, the difference between each dimension at the *n*-th position and the previous (*n* − 1) position can be either 1 or −1 [89]. Therefore, we may define the following set of equations in order to update the values of each dimension array considering that *x*[−1] = *y*[−1] = *z*[−1] = 0:

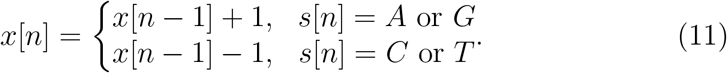

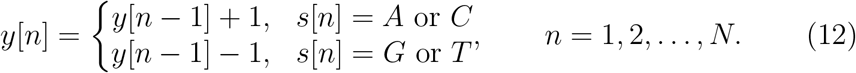

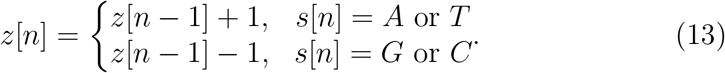

Finally, the DFT and power spectrum may be defined as [94]:

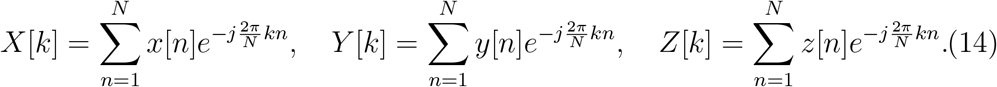

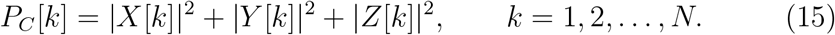

#### 3.3.5. EIIP Representation

Nair and Sreenadhan [90] proposed EIIP values of nucleotides to represent biological sequences and to locate exons. According to the authors, a numerical sequence representing the distribution of free electron energies can be called *“EIIP indicator sequence”*, e.g., let **s** = (G, A, G, A, G, T, G, A, C, C, A), thus, **b** = (0.0806, 0.1260, 0.0806, 0.1260, 0.0806, 0.1335, 0.0806, 0.1260, 0.1340, 0.1340, 0.1260), as shown in Equation (16). The DFT and power spectrum of this representation are presented in Equation (17).

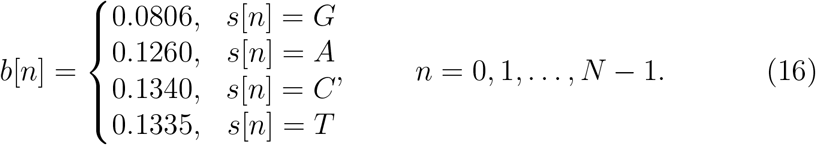

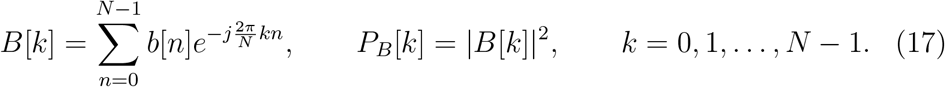

#### 3.3.6. Complex Numbers Representation

This numerical mapping has the advantage of better translating some of the nucleotides features into mathematical properties [92] and represents the complementary nature of AT and CG pairs [84]; e.g., let **s** = (G, A, G, A, G, T, G, A, C, C, A), thus, **r̅** = (−1 − *j*, 1 + *j*, −1 − *j*, 1 + *j*, −1 − *j*, 1 − *j*, −1 − *j*, 1 + *j*, −1 + *j*, −1 + *j*, 1 + *j*), as shown in Equation (18). The DFT and power spectrum of this representation are presented in Equation (19).

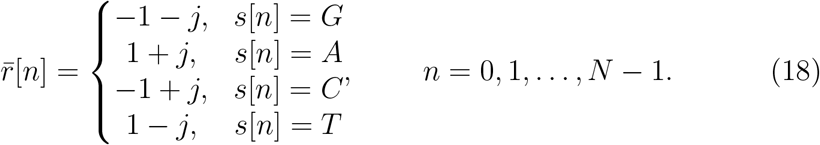

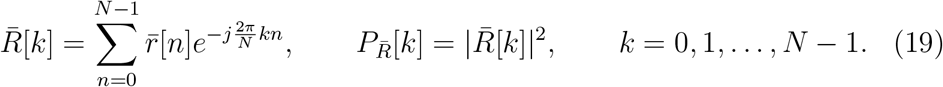

#### 3.3.7. Features

The feature extraction is applied in each representation with Fourier transform, adopting Peak to Average Power Ratio (PAPR), mistakenly confused with the Signal to Noise Ratio (SNR), average power spectrum, median, maximum, minimum, sample standard deviation, population standard deviation, percentile (15/25/50/75), amplitude, variance, interquartile range, semi-interquartile range, coefficient of variation, skewness, and kurtosis. According to [95], the RNA has a statistical phenomenon known as period-3 behavior or 3-base periodicity, where the peak power will always be at the sample *N/*3. The PAPR is defined as [96]:

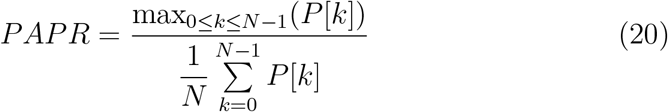

### 3.4. Entropy

Information theory has been widely used in bioinformatics [97, 98]. Based on this, we consider the study of [99], which applied an algorithmic and mathematical approach to DNA code analysis using entropy and phase plane. According to [98], entropy is a measure of the uncertainty associated with a probabilistic experiment. To generate a probabilistic experiment, we use a known approach in bioinformatics, the k-mer (our pipeline - Figure 4).

**Figure 4:**
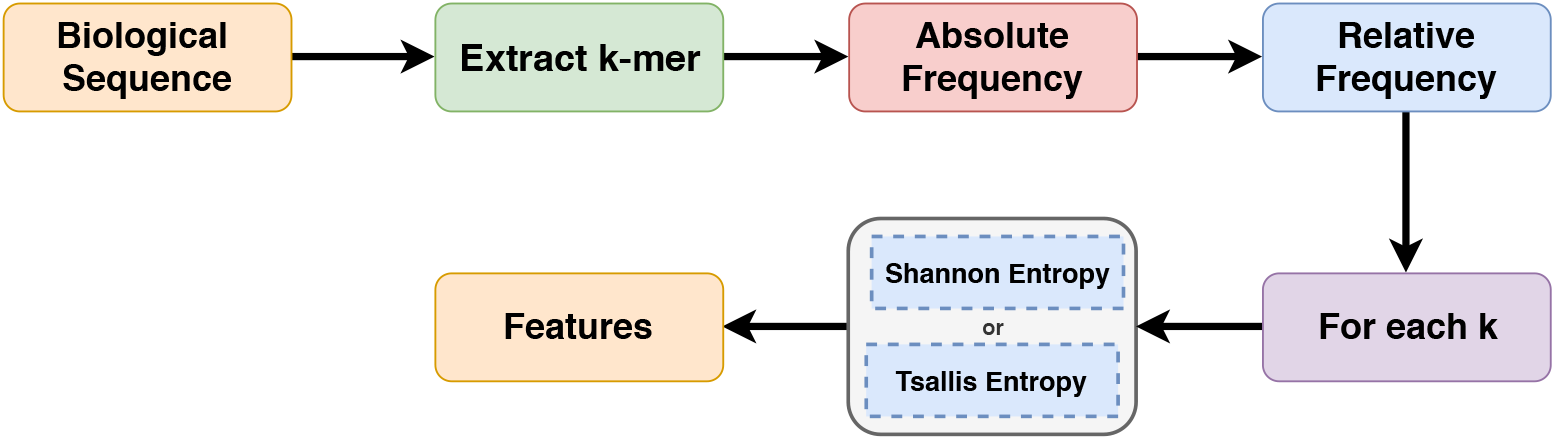
Entropy Pipeline. (1) Each sequence is mapped in *k*-mers; (2) The absolute frequency of each *k* is calculated; (3) Based on absolute frequency, the relative frequency is calculated; (4) The Tsallis or Shannon entropy is applied to each *k*.

In this method, each sequence is mapped in the frequency of neighboring bases k, generating statistical information. The k-mer is denoted by *P_k_*, corresponding to Equation (21).

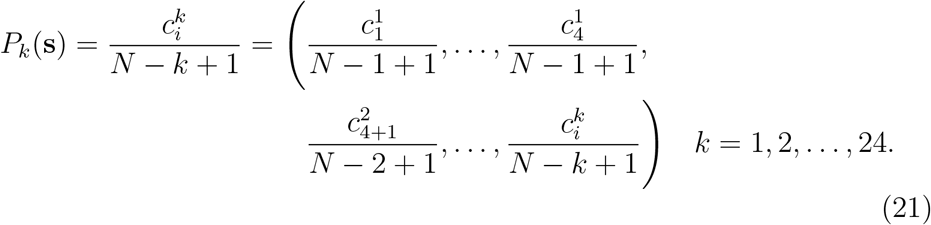

We applied this equation to each sequence with frequencies of *k* = 1, 2, …, 24. Where, 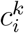 is the number of substring occurrences with length *k* in a sequence (**s**) with length *N*, in which the index *i* ∈ {1, 2, …, 4^1^ + … + 4^*k*^} represents the analyzed substring. For a better understanding, Figure 5 demonstrated an example with *k* = 6 and *k* = 9.

**Figure 5:**
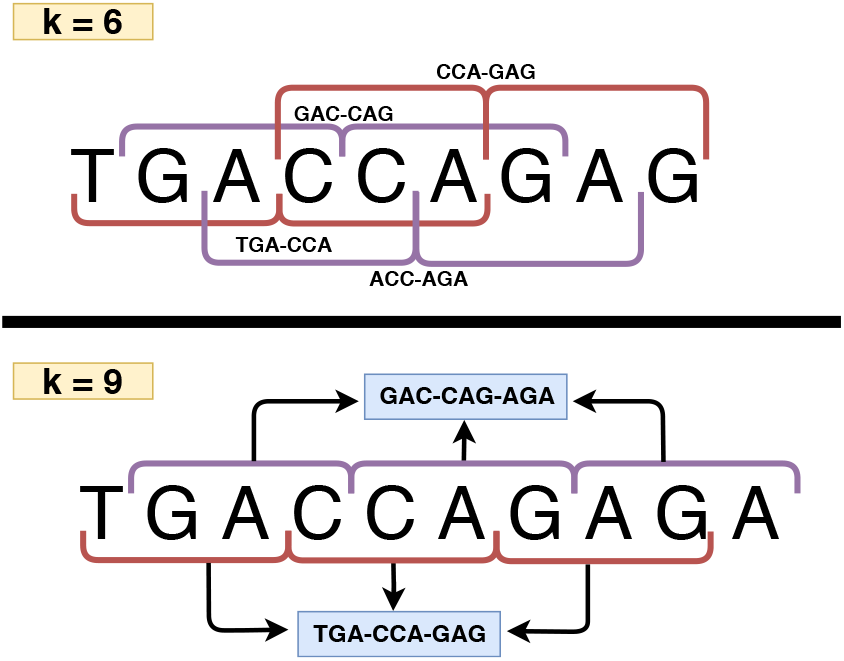
*k*-mer Workflow. Example with *k* = 6 and *k* = 9.

Basically, histograms with short bins are adopted, such as [{*A*}, {*C*}, {*G*}, {*T*}], that occur for *k* = 1, up to histograms with long sequence counting bins such as [{*GGGGGGGGGGGG*}, …, {*AAAAAAAAAAAA*}], that result for *k* = 12. Where, after counting the absolute frequencies of each *k*, we generate relative frequencies (see Equation (21)), and then apply Shannon and Tsallis entropy to generate the features.

#### 3.4.1. Shannon and Tsallis Entropy

Fundamentally, we chose Shannon entropy, because it quantifies the amount of information in a variable [100], that is, we can reach a single value that quantifies the information contained in different observation periods (e.g., our case: k-mer). However, according to [101], it is important to explore a generalized form of the Shannon’s entropy. Based on this, we have opted for a generalized entropy proposed by Tsallis, applied by several works in the literature [102, 103]. Thereby, for a discrete random variable *F* taking values in {*f*[0], *f*[1], *f*[2], …, *f*[*N* − 1]} with probabilities {*p*[0], *p*[1], *p*[2], …, *p*[*N* − 1]}, represented as *P*(*F* = *f*[*n*]) = *p*[*n*]. The Shannon (Equation 22) and Tsallis (Equation 23) entropy associated with this variable is given by the following expressions:

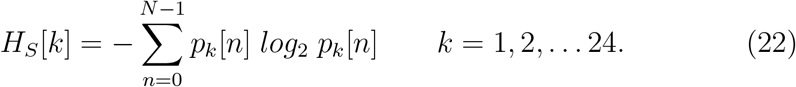

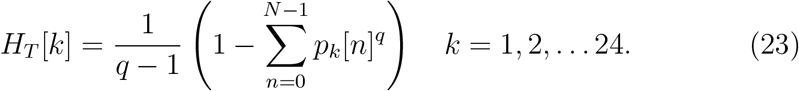

Where *k* represents the analyzed *k*-mer, *N* the number of possible events and *p*[*n*] the probability that event *n* occurs.

### 3.5. Complex Networks

Complex networks are widely used in mathematical modeling and have been an extremely active field in recent years [104], as well as becoming an ideal research area for mathematicians, computer scientists, and biologists. Based on this, we consider the study of [66], in which we propose a feature extraction model based on complex networks, as shown in Figure 6.

**Figure 6:**
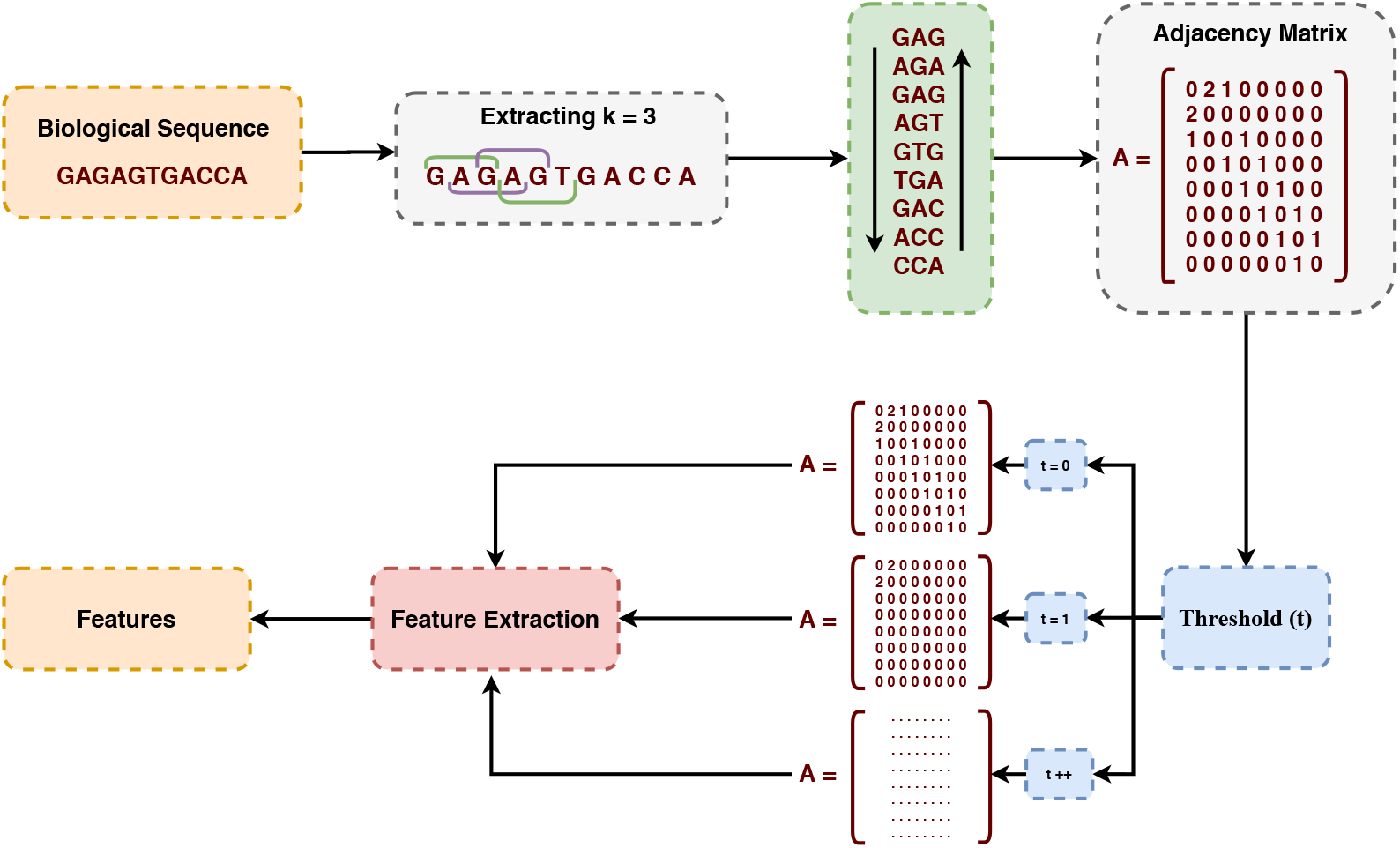
Complex Networks Pipeline. (1) Each sequence is mapped in the frequency of neighboring bases k (k = 3); (2) This mapping is converted to a undirected graph represented by an adjacency matrix; (3) Feature extraction is performed using a threshold scheme; Finally, (4) the features are generated.

Each sequence is mapped to the frequency of neighboring bases k (k = 3 - see Figure 5). This mapping is converted into an undirected graph represented by an adjacency matrix, in which we applied a threshold scheme for feature extraction, thus generating our characteristic vector. Fundamentally, we represent our structure by undirected weighted graphs. According to [104], a graph *G* = {*V, E*} is structured by a set *V* of vertices (or nodes) connected by a set *E* of edges (or links). Each edge reflects a link between two vertices, e.g., *e_p_* = (*i, j*) connection between the vertices *i* and *j* [104]. If there is an edge connecting the vertices *i* and *j*, the elements *a_ij_* are equal to 1, and equal to 0 otherwise.

In our case, the graph is undirected, that is, the adjacency matrix *A* is symmetric, e.g., elements *a_ij_* = *a_ji_* for any *i* and *j* [104]. Furthermore, we apply a threshold scheme presented by [66], in which we extract weight of the edges to capture adjacencies at different frequencies. Finally, as features, several network characterization measures were obtained, based on [66, 105], among them: Betweenness, assortativity, average degree, average path length, minimum degree, maximum degree, degree standard deviation, frequency of motifs (size 3 and 4), clustering coefficient.

### 3.6. Normalization, Training and Evaluation Metrics

Data normalization is a preprocessing technique often applied to a dataset. Essentially, features can have different dynamic ranges. This problem may have a stronger effect in the induction of a predictive model, mainly for distance-based ML algorithms. Consequently, the application of a normalization procedure makes the ranges similar, reducing this problem [106]. We used the min-max normalization, which reduces the data range to 0 and 1 (or −1 to 1, if there are negative values) [20]. The general formula is given as (Equation (24)) [107]:

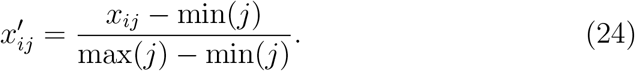

Where *x* is the original value and 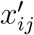 is its normalized version. Furthermore, min(*j*) and max(*j*) are, respectively, the smallest and largest values of 413 a feature *j* [6, 107]. Next, we investigate three classification algorithms, such 414 as ndom Forest (RF) [108], AdaBoost [109] and CatBoost [110]. We chose 415 these ML algorithms because they induce interpretable predictive models 416 when humans can easily understand the internal decision-making process. 417 Thus, domain experts can validate the knowledge used by the models for the classification of new sequences [6]. Finally, to induce our models, we used 70% of samples for *training* (with 10-fold cross-validation) and 30% for *testing*, as shown in Table 2.

**Table 2:** Number of sequences used for training and testing in each dataset.

| Case Study | Dataset | Samples | Training | Testing |
| --- | --- | --- | --- | --- |
| **I** | *A. trichopoda* | 9112 | 6378 | 2734 |
|  | *A. thaliana* | 5080 | 3556 | 1524 |
|  | *C. sinensis* | 4430 | 3101 | 1329 |
|  | *C. sativus* | 3460 | 2422 | 1038 |
|  | *R. communis* | 6974 | 4881 | 2093 |
| **II** | *lncRNA vs. sncRNA* | 2582 | 1807 | 775 |
|  | *lncRNA vs. Antisense* | 114 | 79 | 35 |
|  | *circRNA vs. lncRNA* | 5080 | 3556 | 1524 |

The methods were evaluated with four measures: Sensitivity (SE - Equation 26), Specificity (SPC - Equation 27), Accuracy (ACC - Equation 25), and Cohen’s kappa coefficient [111] (Equation 28).

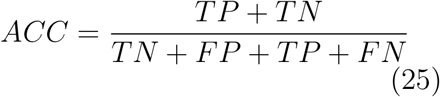

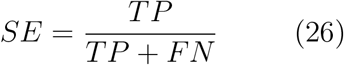

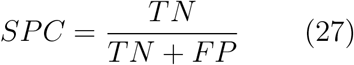

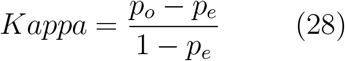

These measures use True Positive (TP), True Negative (TN), False Positive (FP) and False Negative (FN) values, where: TP measures the correctly predicted positive label; TN represents the correctly classified negative label; FP describes all those negative entities that are incorrectly classified as positive and; FN represents the positive label that are incorrectly classified as the negative label.

## 4. Results

This section shows experimental results from 9 feature extraction approaches with mathematical models for biological sequences, divided into two parts: Case Study I and Case Study II.

### 4.1. Case Study I

Initially, we induced models with the RF, AdaBoost, and CatBoost classifiers in the training set of three datasets (*A. trichopoda*, *A. thaliana*, and *R. communis*, we randomly chose three datasets for evaluating the classifiers). Our initial goal is to choose the best classifier to follow in the testing phases. Thereby, to estimate the real accuracy, we applied 10-fold cross-validation, as shown in Table 3.

**Table 3:** Accuracy for the training set (*A. trichopoda*, *A. thaliana*, and *R. communis*) using 10-fold cross-validation.

| Dataset | Model | RF | AdaBoost | CatBoost |
| --- | --- | --- | --- | --- |
| *A. trichopoda* | Z-curve | 0.90 (± 0.03) | 0.91 (± 0.02) | **0.92 (± 0.02)** |
| | Binary | 0.92 (± 0.02) | **0.94 (± 0.02)** | **0.94 (± 0.02)** |
| | Real | 0.91 (± 0.02) | 0.93 (± 0.02) | **0.94 (± 0.02)** |
| | Integer | 0.91 (± 0.02) | 0.93 (± 0.02) | **0.94 (± 0.02)** |
| | EIIP | 0.92 (± 0.02) | **0.94 (± 0.02)** | **0.94 (± 0.02)** |
| | Complex | 0.92 (± 0.03) | **0.94 (± 0.02)** | **0.94 (± 0.02)** |
| | Graphs | 0.92 (± 0.02) | **0.94 (± 0.02)** | **0.94 (± 0.02)** |
| | Shannon | 0.92 (± 0.02) | **0.94 (± 0.02)** | **0.94 (± 0.02)** |
| | Tsallis | 0.92 (± 0.02) | **0.94 (± 0.02)** | **0.94 (± 0.02)** |
| *A. thaliana* | Z-curve | **0.95 (± 0.02)** | 0.93 (± 0.03) | 0.94 (± 0.02) |
|  | Binary | **0.94 (± 0.02)** | **0.94 (± 0.02)** | **0.94 (± 0.02)** |
| | Real | **0.95 (± 0.02)** | 0.94 (± 0.02) | **0.95 (± 0.02)** |
|  | Integer | **0.94 (± 0.02)** | **0.94 (± 0.02)** | **0.94 (± 0.02)** |
| | EIIP | **0.95 (± 0.02)** | 0.94 (± 0.02) | 0.95 (± 0.03) |
| | Complex | 0.94 (± 0.02) | 0.94 (± 0.02) | **0.94 (± 0.01)** |
| | Graphs | 0.94 (± 0.02) | 0.94 (± 0.02) | **0.95 (± 0.02)** |
| | Shannon | 0.94 (± 0.02) | 0.94 (± 0.02) | **0.95 (± 0.02)** |
|  | Tsallis | **0.94 (± 0.02)** | **0.94 (± 0.02)** | **0.94 (± 0.02)** |
| *R. communis* | Z-curve | **0.93 (± 0.02)** | 0.92 (± 0.02) | **0.93 (± 0.02)** |
| | Binary | **0.95 (± 0.01)** | 0.95 (± 0.02) | 0.95 (± 0.02) |
| | Real | **0.95 (± 0.02)** | 0.94 (± 0.02) | 0.94 (± 0.02) |
| | Integer | **0.94 (± 0.01)** | **0.94 (± 0.01)** | 0.94 (± 0.02) |
| | EIIP | 0.95 (± 0.02) | 0.95 (± 0.02) | **0.95 (± 0.01)** |
| | Complex | 0.95 (± 0.02) | **0.95 (± 0.01)** | **0.95 (± 0.01)** |
| | Graphs | **0.95 (± 0.01)** | **0.95 (± 0.01)** | 0.95 (± 0.02) |
| | Shannon | 0.95 (± 0.02) | 0.95 (± 0.02) | **0.95 (± 0.01)** |
|  | Tsallis | **0.95 (± 0.01)** | **0.95 (± 0.01)** | **0.95 (± 0.01)** |

Assessing each classifier, we noted that the best performance was of the CatBoost with all mathematical models in *A. trichopoda*, followed by AdaBoost (6 best results) and RF (no better results). In *A. thaliana*, CatBoost kept the best performance (7 best results), followed by RF (6 best results) and AdaBoost (3 best results). In contrast, the RF classifier obtained the best results (6) in *R. communis*, followed by CatBoost (5 best results) and AdaBoost (3 best results). Based on this, we continued testing the models with the CatBoost classifier. Thus, in Table 4, we present the results of all mathematical models using 4 evaluation metrics.

**Table 4:**
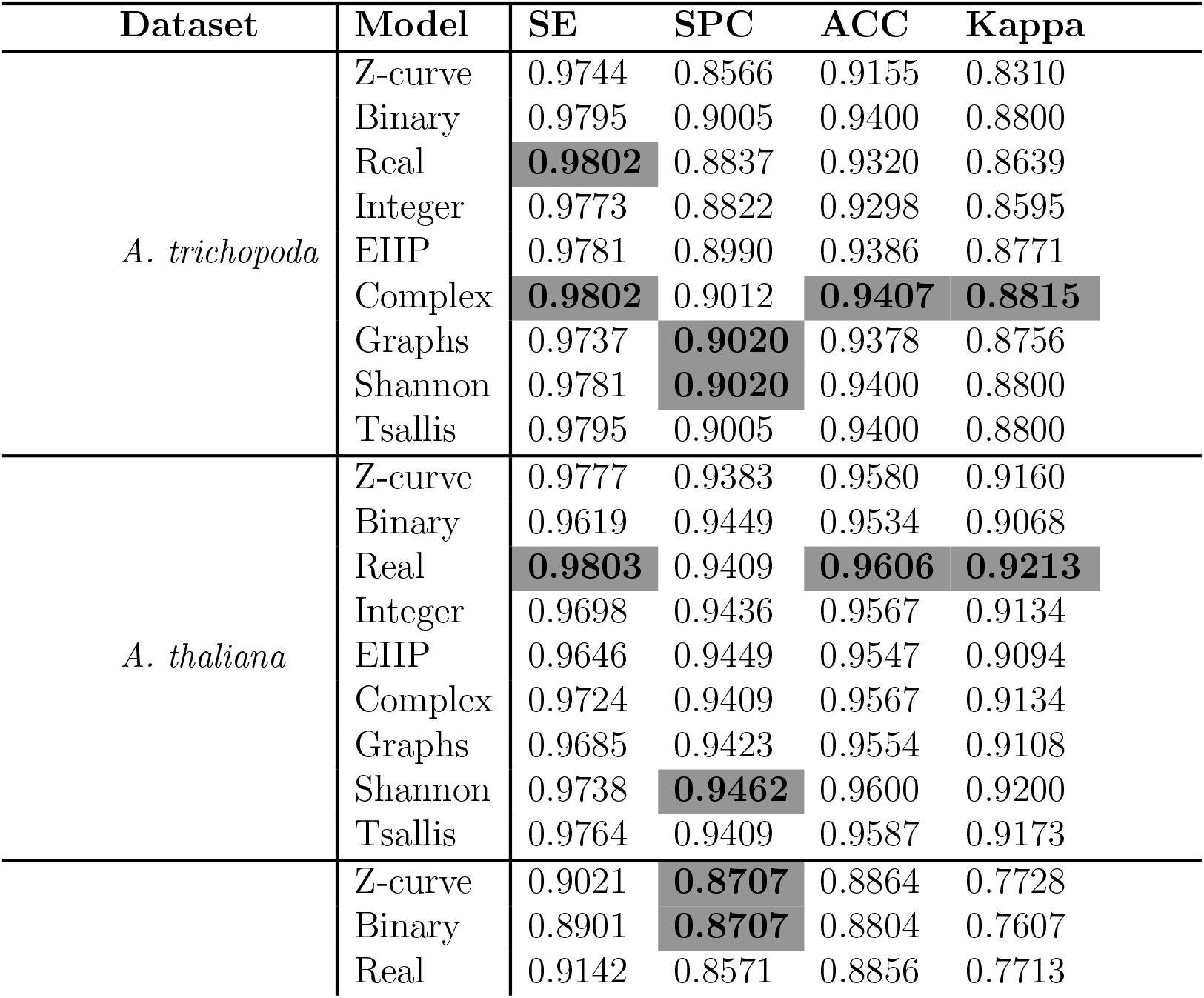
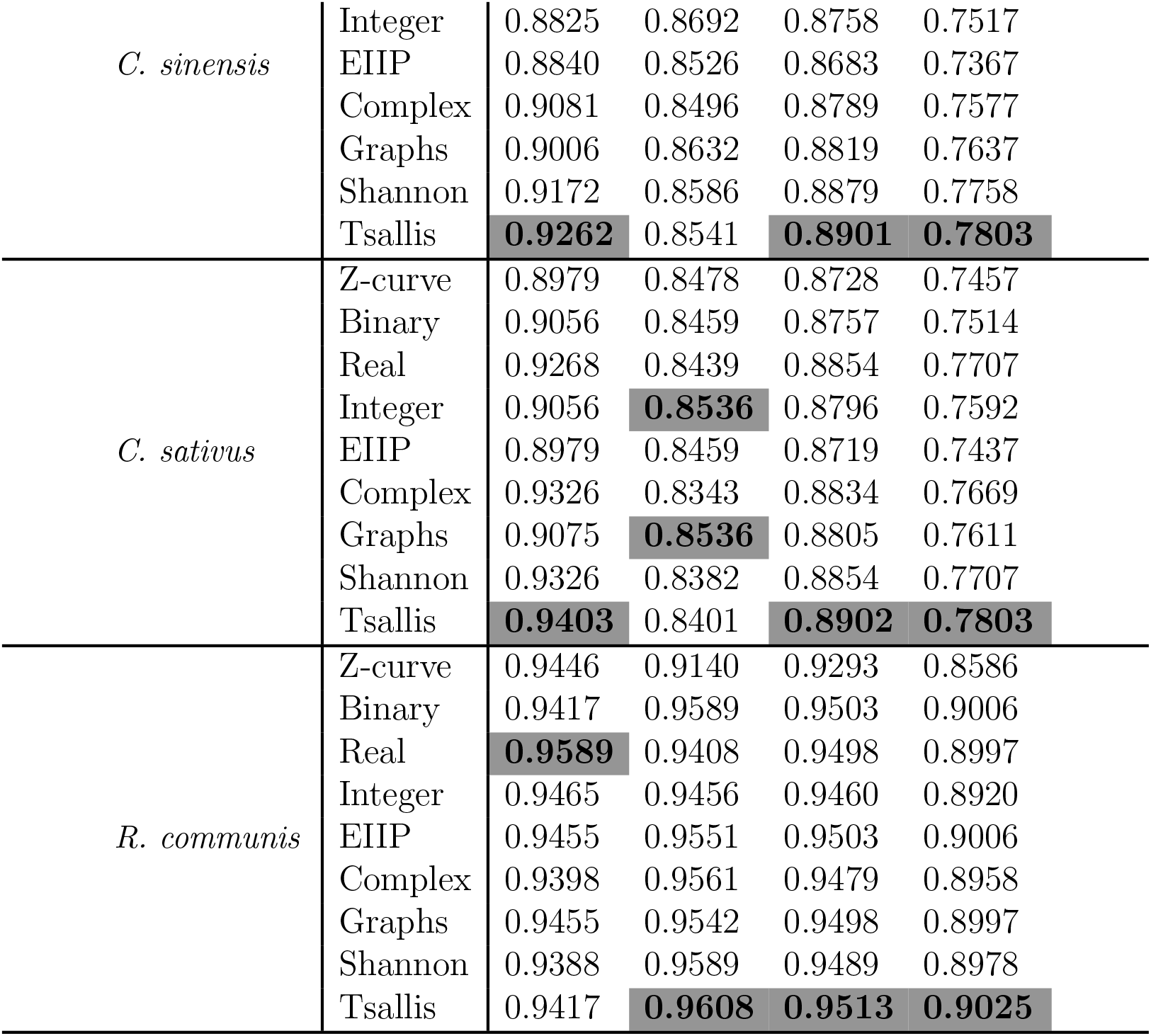
Performance analysis. This table compares the sensitivity, specificity, accuracy and kappa metrics for each model in the test sets using CatBoost classifier.

As can be seen, all models presented robust results, with the worst performance (ACC) of 0.8901 (*C. sinensis*) and the best of 0.9606 (*A. thaliana*). That is, all models were robust in different datasets without a high loss of performance. Assessing each metric individually, we realized that in SE, the best performance was from Real representation (3 datasets), followed by Tsallis (2 datasets) and Complex numbers (1 dataset). In SPC, the best results were from Entropy (3 datasets), followed by Graphs (2 datasets). In ACC, Tsallis presented the best performance (3 datasets), followed by Real representation and Complex numbers (1 dataset). For each dataset, we can see in *A. trichopoda* the best ACC was 0.9407 (Complex); *A. thaliana* with 0.9606 (Real); *C. sinensis* with 0.8901 (Tsallis); *C. sativus* with 0.8902 (Tsallis); and *R. communis* with 0.9513 (Tsallis). Highlight for Tsallis entropy, which presented the best results, mainly in accuracy, proving to be more generalist in the case study I.

### 4.2. Case Study II

After evaluating all methods in 5 datasets (lncRNA of different species) and observing their results, we applied a second case study, where we used only three mathematical models for generalization analysis, including GSP (Fourier + complex numbers), entropy (Tsallis) and graphs (complex networks). Here, our objective was to analyze how each model (feature extraction approach) behaved in different biological sequence classification problems. In other words, we assessed the generalization of each approach to classifying sequences with different structures (distinct problem). For this, we tested 3 new datasets established in Section 3.1.2, as can be seen in Figure 7.

**Figure 7:**
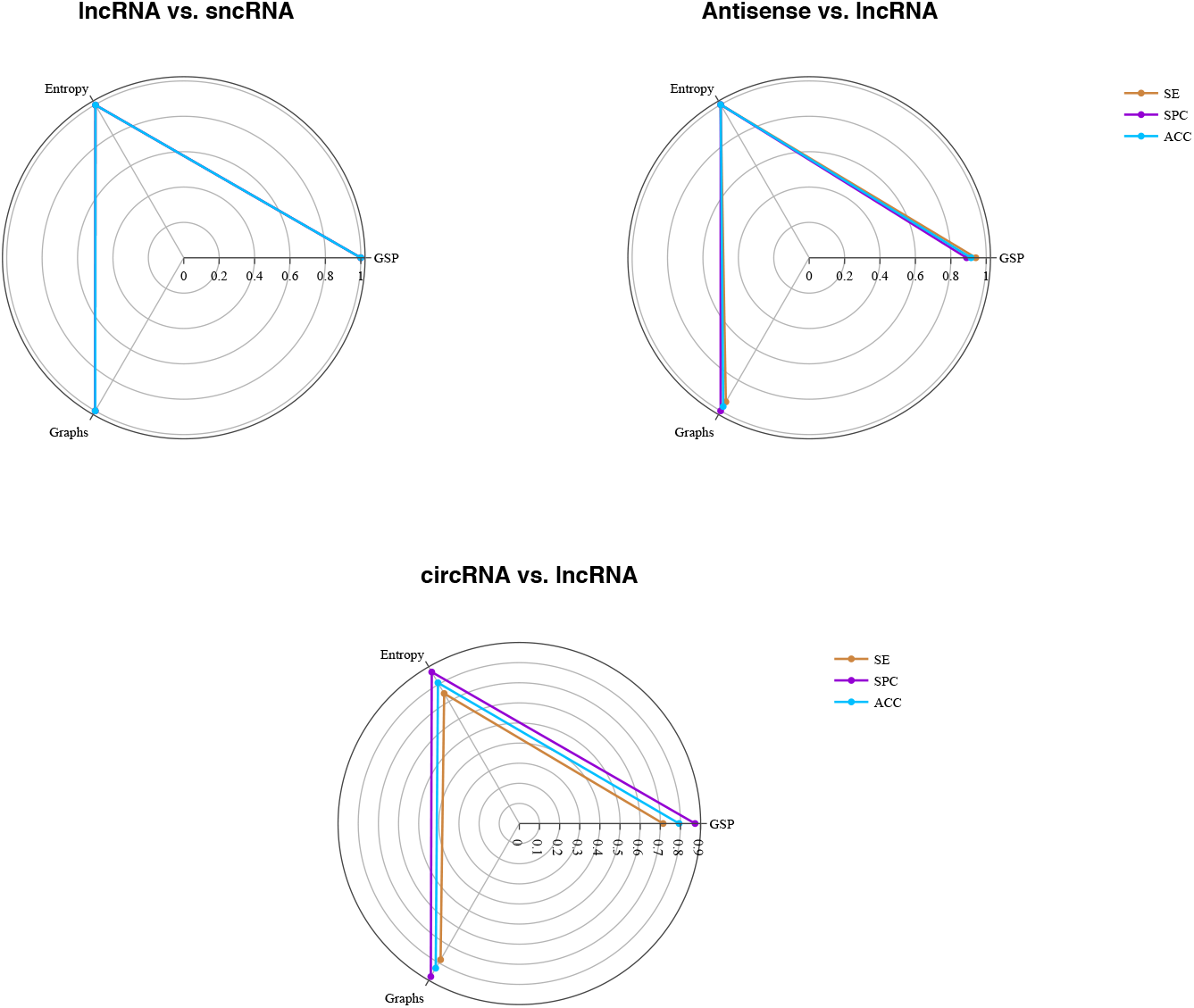
Performance analysis of three mathematical models, GSP (fourier + complex numbers), entropy (Tsallis) and graphs (complex networks), for different problems.

Again, all showed robust results, in which, graph-based models are the best in 2 of the 3 problems analyzed, followed by entropy and GSP. In the three datasets, our approaches have achieved relevant results with ACC, SE and SPC, proving to be efficient and generalist, when exposed to different problem scenarios. Furthermore, if we analyze at the last problem (circRNA vs. lncRNA), our approaches were effective when compared to our references that reached an ACC of 0.7780 [21] and 0.7890 [27] in their datasets against 0.8307 from our best model (graph - using these comparisons as an (indirect) reference indicator).

### 4.3. Statistical Significance Tests

The statistical significance was assessed in both case studies (difference in ACC), using Friedman’s statistical test and the Conover post-hoc test. Thereby, our null hypothesis (*H*0 = *M* (1) = *M* (2) = *…* = *M* (*k*)), is tested against the alternative hypothesis (*H_A_* = at least one model has statistical significance (*α* = 0.05, *p < α*)). First, we apply the global test in the case study I, in which the Friedman test indicates significance (*χ*^2^(8) = 17.34, *p*-value = 0.0268), that is, we can reject *H*0, as *p* < 0.05. Thus, it is essential to execute the post-hoc statistical test. Conover statistics values were obtained, as well as *p*-values (see Table 5), using 95% of significance (*α* = 0.05).

**Table 5:** Conover statistics values - The accepted alternative hypothesis is in bold (*p*-values for *α* = 0.05).

|  | Z-curve | Binary | Real | Integer | EIIP | Complex | Graphs | Shannon |
| --- | --- | --- | --- | --- | --- | --- | --- | --- |
| Binary | 0.5580 | - | - | - | - | - | - | - |
| Real | 0.1416 | 0.3671 | - | - | - | - | - | - |
| Integer | 0.7896 | 0.3956 | 0.0852 | - | - | - | - | - |
| EIIP | 0.9574 | 0.5230 | 0.1284 | 0.8309 | - | - | - | - |
| Complex | 0.3671 | 0.7489 | 0.5580 | 0.2451 | 0.3399 | - | - | - |
| Graphs | 0.5580 | 1.0000 | 0.3671 | 0.3956 | 0.5230 | 0.7489 | - | - |
| Shannon | 0.0687 | 0.2057 | 0.7089 | **0.0390** | 0.0616 | 0.3399 | 0.2057 | - |
| Tsallis | **0.0146** | 0.0550 | 0.2898 | **0.0075** | **0.0128** | 0.1050 | 0.0550 | 0.4892 |

Concerning to the Conover post-hoc test, entropy-based models have highly significant differences for the Z-curve (*p* < 0.0146), Integer (*p* < 0.0075 - Tsallis and *p* < 0.0390 - Shannon), and EIIP (*p* < 0.0128). Possibly, these results indicate that entropy has a more significant performance when compared to representations with Fourier. However, other mathematical models in case study I do not differ significantly, indicating their efficiency in all datasets. Now, evaluating case study II, we realized that the global test with Friedman’s statistical test is not significant, in which we obtained *χ*^2^(2) = 1.64, *p*-value = 0.4412, indicating that the three studied feature extraction techniques show a similar performance in the problems, once more confirming the effectiveness and robustness of all mathematical models.

### 4.4. Computational Time

In addition, we also assessed the computational time cost of each tested model. To do this, we ran three models, GSP (Fourier + complex numbers), entropy (Tsallis) and graphs (complex networks)), in 1291 random sequences, as shown in Figure 8.

**Figure 8:**
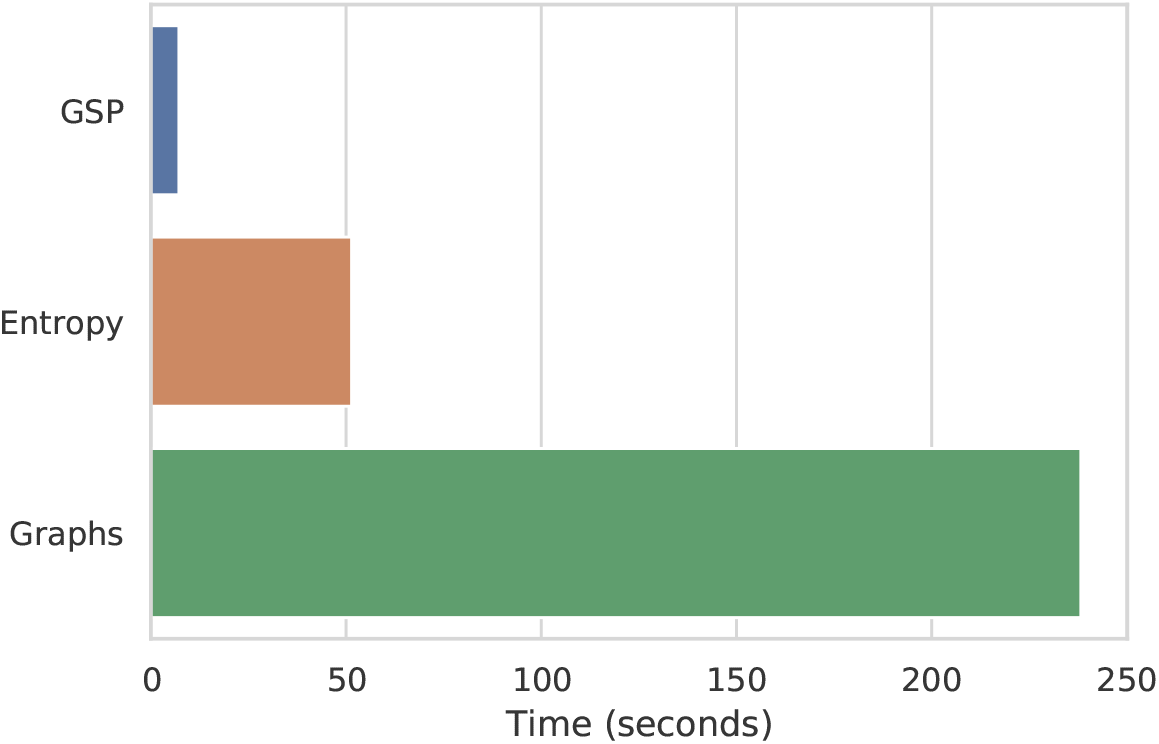
Execution Time.

We performed the experiments using Intel Core i3-9100F CPU (3.60GHz), 16GB memory, and running in Debian GNU/Linux 10. The lowest cost in computational time is for models based on GSP (0m7.183s) and entropy (0m51.427s), while graphs (3m58.208s) have a much higher cost. These results demonstrated that, although the models present a similar performance, the computational time efficiency is significantly different.

## 5. Discussion

This section discusses our findings in terms of whether they support our hypothesis (*feature extraction approaches based on mathematical models are as efficient and generalist as biological approaches*). Overall, several experimental tests were assumed in this research, in which all feature extraction approaches based on mathematical models showed excellent results, as can be seen in Table 4 and Figure 7. Regarding its performance in distinct classification problems, case study II, we used only three mathematical models for generalization analysis, including GSP (Fourier + complex numbers), entropy (Tsallis) and graphs (complex networks). In which, entropy and graphbased models reported the best performance followed by GSP. Furthermore, all models maintained robust results in different sequence classification problems.

Furthermore, to fully support our hypothesis, we also compare three mathematical models shown in Figure 7 concerning a biological and hybrid approach, in four datasets ((lncRNA vs. mRNA (case study I)); (lncRNA vs. sncRNA; lncRNA vs. Antisense; circRNA vs. lncRNA (case study II)). Thus, we generate our biological model using some of the most applied features in Figure 1. Thus, features used by the models are:

- **Biological:** The features were provided by [25]: Fickett TESTCODE score, isoelectric point, open reading frame (ORF) length, and ORF integrity.
- **Hybrid:** The features were generated by one of the most current approaches in the literature (lncFinder [26] - 2018). We classify this model as a hybrid because it uses a combination of biological and mathematical features. Among the biological characteristics is Logarithm-distance of hexamer on ORF, length and coverage of the longest ORF. Regarding mathematical features, [26] uses an EIIP-based physicochemical property with Fourier Transform (similar to our approach with GSP, but using only EIIP mapping).

For a fair comparison, the new experiments follow the same methodology (70% training, 30% test, and CatBoost classifier), as shown in Table 6.

**Table 6:**
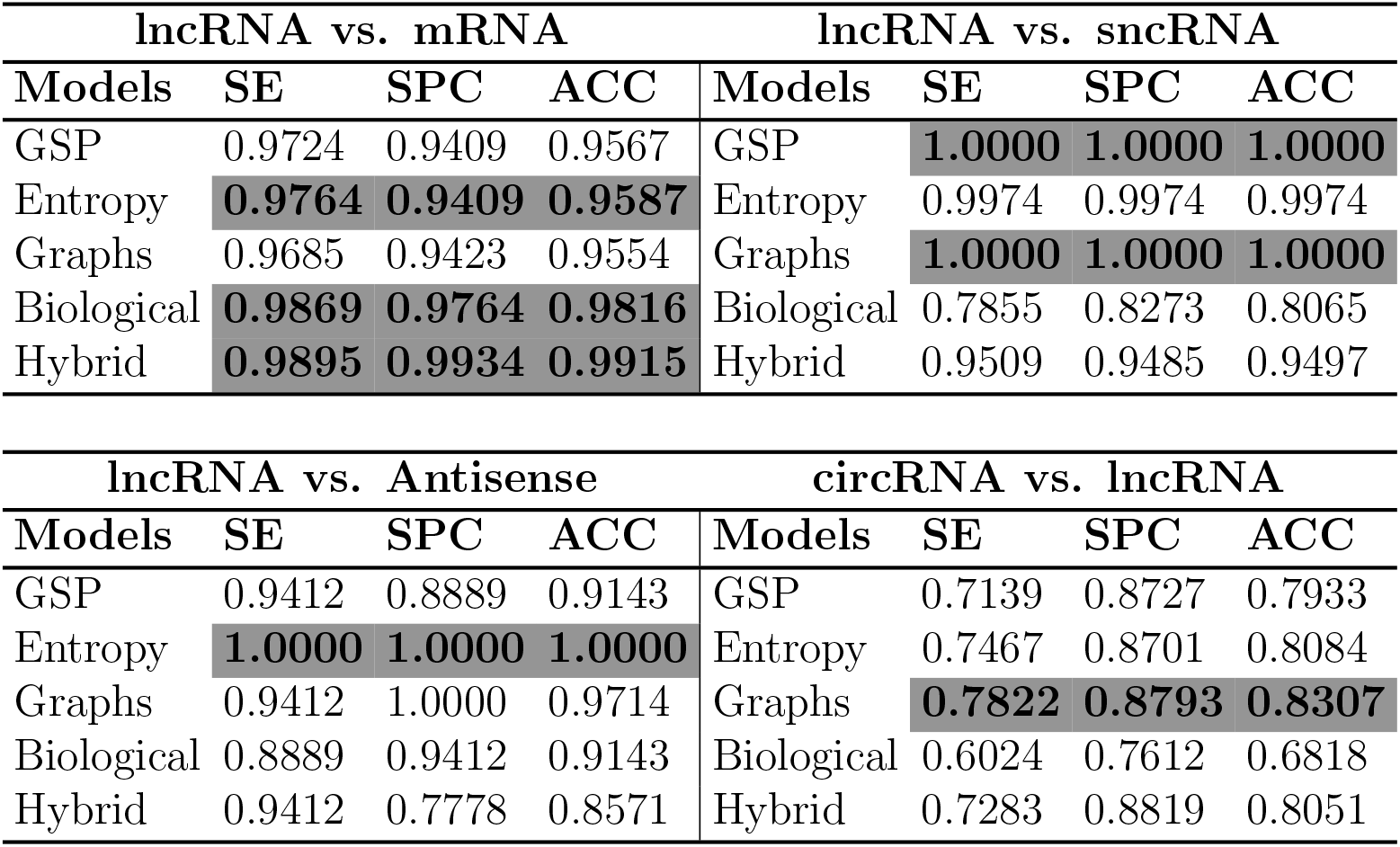
Performance analysis of three mathematical models against a biological and hybrid model for different sequence classification problems.

As can be seen, the hybrid model (0.9915) reported the best performance in the first dataset (lncRNA vs. mRNA), followed by the biological (0.9816) and our mathematical model (Entropy - 0.9587), with only a difference of 0.0328 and 0.0229, respectively. However, it is relevant to highlight that the biological and hybrid models use the ORF descriptor, a highly employed feature for discovering coding sequences and which, according to [19] is an essential guideline for distinguishing lncRNAs from mRNA. In other words, this explains the great result, but, as mentioned at the beginning of this manuscript, this type of feature with a biological insight is often difficult to reuse or adapt to another specific problem. Thereby, our study has an gain in terms of generalization, since this would not be possible only with the ORF. If we analyze at the hybrid model, in this first dataset, the gain was minimal compared to the biological (0.0099), confirming the efficiency of the previously mentioned features (ORF). This is different from our approaches, which showed an robust result without using bias features for the analyzed problem.

Hence, this hypothesis is proven in the other three datasets, where our athematical models perform much better than the biological model, mainly in the fourth dataset (circRNA vs. lncRNA), in which we obtained a gain of 0.1489 in ACC. Regarding the hybrid model, it can be observed that the mixture of biological and mathematical characteristics helped to keep the model competitive in all datasets, indicating the effectiveness of mathematical features. Even so, our models showed the best results in three of the four proposed problems. Therefore, our pipeline is efficient in terms of generalization to classify lncRNA from mRNA, as well as other biological sequence classification problems. We also assessed the statistical significance of the mathematical versus biological approach in the previously applied tests, in which entropy (*p* < 0.0480) and graphs (*p* < 0.0200) indicated significant results concerning the biological model. Lastly, considering all these findings, we fully support the suggested hypothesis.

## 6. Conclusion

This work proposed to analyze feature extraction approaches for biological sequence classification. Specifically, we concentrated our work on the study of feature extraction techniques using mathematical models. We analyzed mathematical models to propose efficient and generalist techniques for different problems. As a case study, we used lncRNA sequences. Moreover, we divided this paper into two case studies. In our experiments, as a starting point, 9 mathematical models for feature extraction were analyzed: 6 numerical mapping techniques with Fourier transform; Tsallis and Shannon entropy; Graphs (complex networks). Thereby, several biological sequence classification problems were adopted to validate the proposed approach.

In our experiments, all mathematical models presented relevant and robust results, with performances (ACC) between 0.8901-0.9606 in case study I. In case study II, once more, all showed effective results with models based on entropy and graphs showing the best performance, followed by GSP. Furthermore, to validate our study, we compared three mathematical models against a biological and hybrid approach, in four different datasets. In which, our models demonstrated suitable results, and was superior or competitive and robust in terms of generalization. Moreover, we verified that mathematical approaches perform as accurately as biological approaches and have a better generalization capacity since they outperform biological features in scenarios not designed for them. Finally, among the different feature extraction approaches tested in this work, the combination of k-mer and entropy, as well as complex networks performs better than GSP at the cost of a significant increase in computational complexity.

## Acknowledgements

The authors would like to thank UTFPR-CP, ICMC-USP, and CAPES for the financial support given to this research.

